# Driver pattern identification over the gene co-expression of drug response in ovarian cancer by integrating high throughput genomics data

**DOI:** 10.1101/145268

**Authors:** Xinguo Lu, Jibo Lu, Bo Liao, Keqin Li

## Abstract

The multiple types of high throughput genomics data create a potential opportunity to identify driver pattern in ovarian cancer, which will acquire some novel and clinical biomarkers for appropriate diagnosis and treatment to cancer patients. However, it is a great challenging work to integrate omics data, including somatic mutations, Copy Number Variations (CNVs) and gene expression profiles, to distinguish interactions and regulations which are hidden in drug response dataset of ovarian cancer. To distinguish the candidate driver genes and the corresponding driving pattern for resistant and sensitive tumor from the heterogeneous data, we combined gene co-expression modules and mutation modulators and proposed the identification driver patterns method. Firstly, co-expression network analysis is applied to explore gene modules for gene expression profiles via weighted correlation network analysis (WGCNA). Secondly, mutation matrix is generated by integrating the CNVs and somatic mutations, and a mutation network is constructed from this mutation matrix. The candidate modulators are selected from the significant genes by clustering the vertex of the mutation network. At last, regression tree model is utilized for module networks learning in which the achieved gene modules and candidate modulators are trained for the driving pattern identification and modulator regulatory exploring. Many of the candidate modulators identified are known to be involved in biological meaningful processes associated with ovarian cancer, which can be regard as potential driver genes, such as CCL11, CCL16, CCL18, CCL23, CCL8, CCL5, APOB, BRCA1, SLC18A1, FGF22, GADD45B, GNA15, GNA11 and so on, which can help to facilitate the discovery of biomarkers, molecular diagnostics, and drug discovery.

---

Ovarian cancer is known as a complex disease of the genome. During tumorigenesis, many factors including genomic affection and epigenomic affection contribute to pathological gene expression changes. Hence, acquiring the etiopathogenesis and chemoresponse of cancer faces a great challenge. And the achievement can be utilized to deploy the high-performance diagnosis and treatment of cancer patients. In spite of many studies presented for ovarian cancer prognosis, currently, there is no completely validated clinical model for predicting ovarian cancer prognosis and drug response. Therefore, it remains an important research issue to identify prognostic and predictive driver patterns for improving ovarian cancer treatment.

Cancer genomes possess a large number of aberrations including somatic mutations and copy number variations (CNVs). Many mutations contribute to cancer progression from the normal to the malignant state. The researches have shown that some aberrations are vital for tumorigenesis and most cancers are caused by a small number of driver mutations developed over the course of about two decades (Cheng et al. 2015; Vogelstein et al. 2013). The detection of these mutations with exceptionally high association between the copy number variations, somatic mutations and gene expression can ascertain disease candidate genes and potential cancer mechanisms. Several large scale cancer genomics projects, such as the Genomic Data Commons (GDC), The Cancer Genome Atlas (TCGA), and International Cancer Genome Consortium (ICGC), etc., have produced a large volume of data and provided us opportunity to integrate different level of gene expression data to identify candidate driver genes and driver pathways and better understand the cancer at the molecular level.

During exploring the large volume of genomics abnormalities generated from large-scale cancer projects, many computational and statistical methods have been proposed to search for this mutation driver patterns. Adib et al. predicted the potential specificity of several putative biomarkers by using gene expression microarray profile to examine their expression in a panel of epithelial tissues and tumors (Adib et al. 2004). Youn et al. proposed a new method to identify cancer driver genes by accounting for the functional impact of mutations on proteins (Youn et al. 2010). Tomasetti et al. combined conventional epidemiologic studies and genome-wide sequencing data to infer a number of driver mutations which is essential for cancer development (Tomasetti et al. 2014). Jang et al. identified differentially expressed proteins involved in stomach cancer carcinogenesis through analyzing comparative proteomes between characteristic alterations of human stomach adenocarcinoma tissue and paired surrounding normal tissue (Jang et al. 2004). Xiong et al. proposed a general framework in which the feature (gene) selection was incorporated into pattern recognition to identify biomarkers (Xiong et al. 2001). Jung et al. identified novel biomarkers of cancer by combining bioinformatics analysis on gene expression data and validation experiments using patient samples and explored the potential connection between these markers and the established oncogenes (Jung et al. 2011). Logsdon et al. identified a new category of candidate tumor drivers in cancer genome evolution which can be regarded as the selected expression regulators (SERs)-genes driving deregulated transcriptional programs in cancer evolution, and uncovered a previously unknown connection between cancer expression variation and driver events by using a novel sparse regression technique (Logsdon et al. 2015). A cancer genome embodies thousands of genomics abnormalities such as single nucleotide variants, large segment variations, structural aberrations and somatic mutations (Chan et al. 2012; Ding et al. 2015). The corresponding malignant state reflects aberrant copy numbers (CN) and/or mutations and the expression patterns in which the mutation and copy number patterns are embedded (Wei et al. 2016). Integration of copy number/mutation data and gene expression data was proposed to discover driver genes by quantifying the impact of these aberrations on the transcriptional changes (Zhang et al. 2016; Kumar et al. 2016; Alderton et al. 2011). The available implementations for the integrative analysis of genomics data includes regression methods, correlation methods and module network methods (Huang et al. 2011).

Driver patterns, including driver genes, driver mutations, driver pathways and core modules, which are considered as cancer biomarkers, are supposed to promote the cancer progression. Thus, identification of driver patterns is vital for providing insights into carcinogenic mechanism. The integration of gene expression, copy number and somatic mutations data to identify genomics alterations that induce changes in the expression levels of the associated genes becomes a common task in cancer mechanism and drug response studies. However, there is still a tough work to integrate information across the different heterogeneous omics data and distinguish driver patterns which can promote the cancer cell to propagate infinitely. Hence, we proposed a driver pattern identification method over the gene co-expression of drug response in ovarian by integrating high throughput genomics data. In the method, we integrated different level genomic data including gene expression, copy number and somatic mutation to identify drivers of resistance and sensitivity to anti-cancer drugs. First, weighted correlation network analysis is applied to acquire the co-expression gene modules. These modules are utilized as initial inputs for the following module networks learning. Then, mutation network is constructed from the mutation matrix generated by integrating the CNVs and somatic mutations. The candidate modulators are selected from the clusters of the vertex of the mutation network. Finally, an optimization model is used to identify the driving pattern and explore the modulator regulatory. We used publicly available and clinically annotated gene expression, CNVs, and somatic mutation datasets from TCGA. The experimental results show that in the driver patterns many of the identified candidate modulators are known to be involved in biological meaningful processes associated with ovarian cancer, which can be regard as potential driver genes.

## Results

We conducted the driver patterns identification and the driving regulatory analysis as following procedure. Firstly, co-expression network were constructed and the gene modules for gene expression profiles were explored. Then, CNVs and somatic mutations were integrated to build the mutation network. The candidate modulators were selected from the clusters of the vertex of network. A regulation procedure was explored for candidate modulators and gene modules using network learning combined with regression tree model. At last, local polynomial regression fitting model was applied to conduct dosage-sensitive and dosage-resistant analysis.

### Data preprocessing

To select a subset of genes whose aberration/expression profiles were significantly different between sensitive and resistant tumors, we applied differential analyses to the gene expression, and CNV data of the two groups. For differential gene expression analysis and to rank CNV genes, we used the EMD approach: EMDomics, which is designed especially for heterogeneous data. We selected genes with a q-value<0.1. For CNV data, first, we mapped genes to the CNV regions to obtain CNV genes; then, we used the EMD approach for the differential analysis and ranking of the CNV genes. We also calculated the frequency of amplification and deletion for each gene in the two groups and selected genes for which the difference between their frequencies is more than 20%. We used a threshold of log2 copy number ratio of 0.3/-0.3 to call amplified/deleted genes. For somatic mutation data, we also calculated the frequency of mutations for all genes across the samples in each groups and selected the genes that were mutated in more than 2% of the tumors.

### Gene modules with co-expression patterns

After the data preprocessing, 2690 genes were selected for co-expression gene modules exploring. We constructed a weighted correlation network and identified the co-expression modules, which was conducted using WGCNA (Osterhoff et al. 2014; Liu et al. 2016), associated with cis-platinum sensitive and resistant. The gene co-expression modules and the corresponding hierarchical clustering dendrograms of these genes are shown in Fig. 1. The color bars correspond to the clusters of genes which can be seen as gene module. We identified thirty-nine modules which are described in thirty-nine colors as turquoise, blue, brown, yellow, green and so on. The top 10 co-expression modules are shown in Table 1. The first row represents the color of the modules, and the second row represents the number of genes in the corresponding module. The rest genes are in grey. These grey genes are not clustered into any modules. The modules list is shown in Additional file 5.

**Figure 1.**
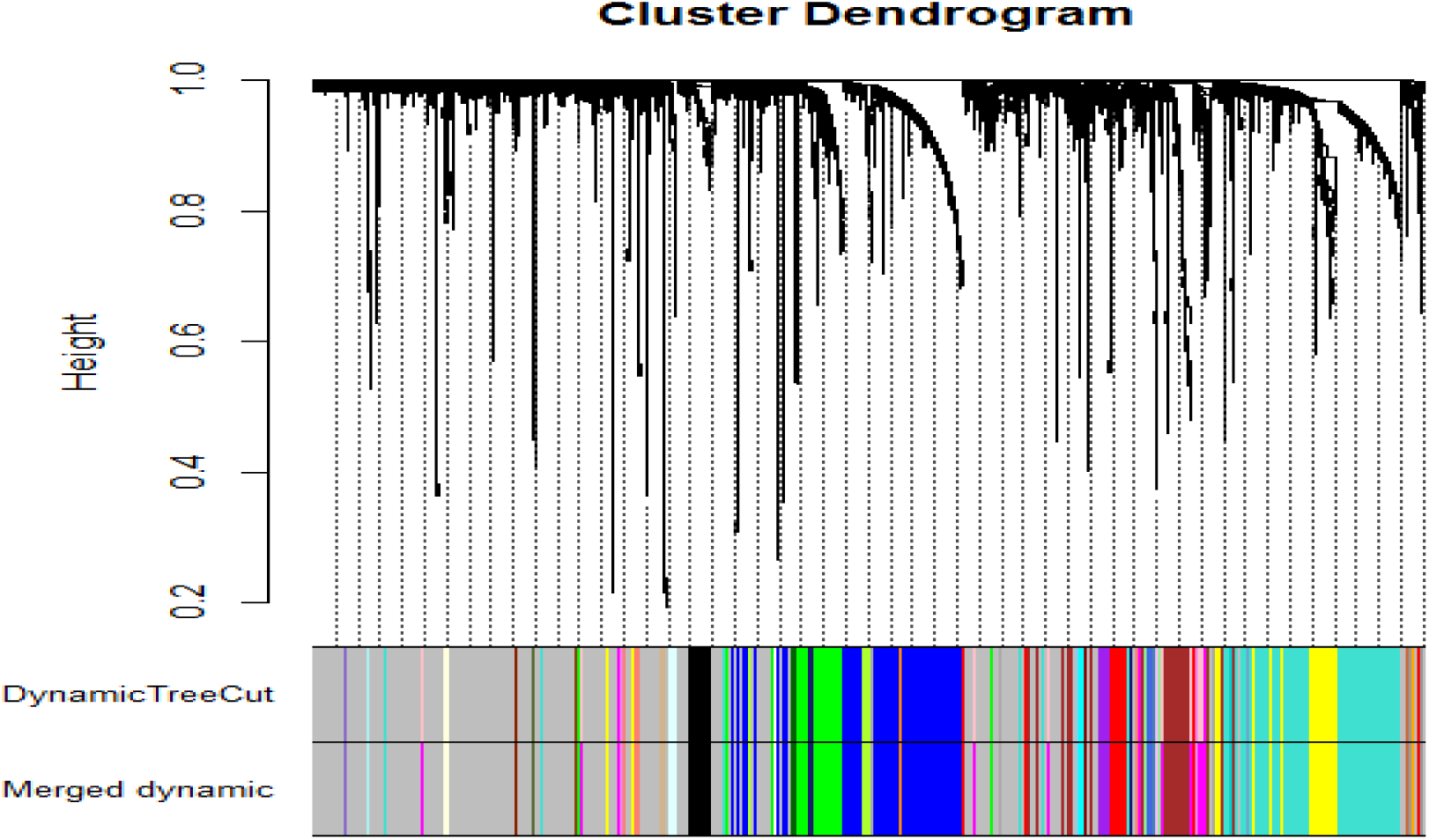
Gene co-expression modules for gene expression profiles. A network heatmap plot (interconnectivity plot) of a gene network together with the corresponding hierarchical clustering dendrograms, with dissimilarity based on topological overlap, together with assigned module colors.

**Table 1.** The top 10 co-expression modules

| module color | turquoise | blue | brown | yellow | green | magenta | red | black | purple | greenyellow |
| --- | --- | --- | --- | --- | --- | --- | --- | --- | --- | --- |
| gene counts | 357 | 288 | 145 | 130 | 115 | 78 | 62 | 61 | 29 | 23 |

### Candidate modulators selected CNVs and somatic mutations

For CNVs data, the mapping of CNVs can promote the search for causal links between genetic variation and disease susceptibility. Thus we mapped gene expression to CNVs regions to obtain CNVs genes, for a specific gene in a given sample, the value of amplified is assigned the number 1, the deleted is assigned the number -1, the normal is assigned the number 0; for somatic mutation data, we first obtained mutated genes and mapped the correlations between somatic mutations and their residing genes. Then, a mutation matrix A is generated by integrating the CNVs and somatic mutations. The mutation matrix is binary: if any variation region in a given gene of the particular sample is in a statistically significant variation or any mutation arises in the specific gene in the given sample, the value of mutation is assigned the number 1; otherwise the value is assigned the number 0. The matrix rows and columns correspond to samples and genes, respectively. A mutation network (MN) was constructed according to the mutation matrix. For a mutated gene, m*_i_* describes the number of mutations in gene i across the samples in the mutation matrix, i.e. 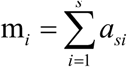. The network vertex weight is defined as *h_i_ =m_i_/m,* where m is the number of samples. Also the edge weight *v_ij_* is defined as the number of samples that exactly one of the pair is mutated divided by the number of samples that at least one of the pair is mutated.

We selected a subgroup of significant genes by clustering the vertex of the mutation network. These genes can be regarded as candidate modulators. We identify a list of 624 initial candidate modulators for the mutation genes (Additional file 6). These modulators includes 40 somatic mutations. The gene expression heatmap of the top 50 candidate modulators is shown in Fig. 2.

**Figure 2.**
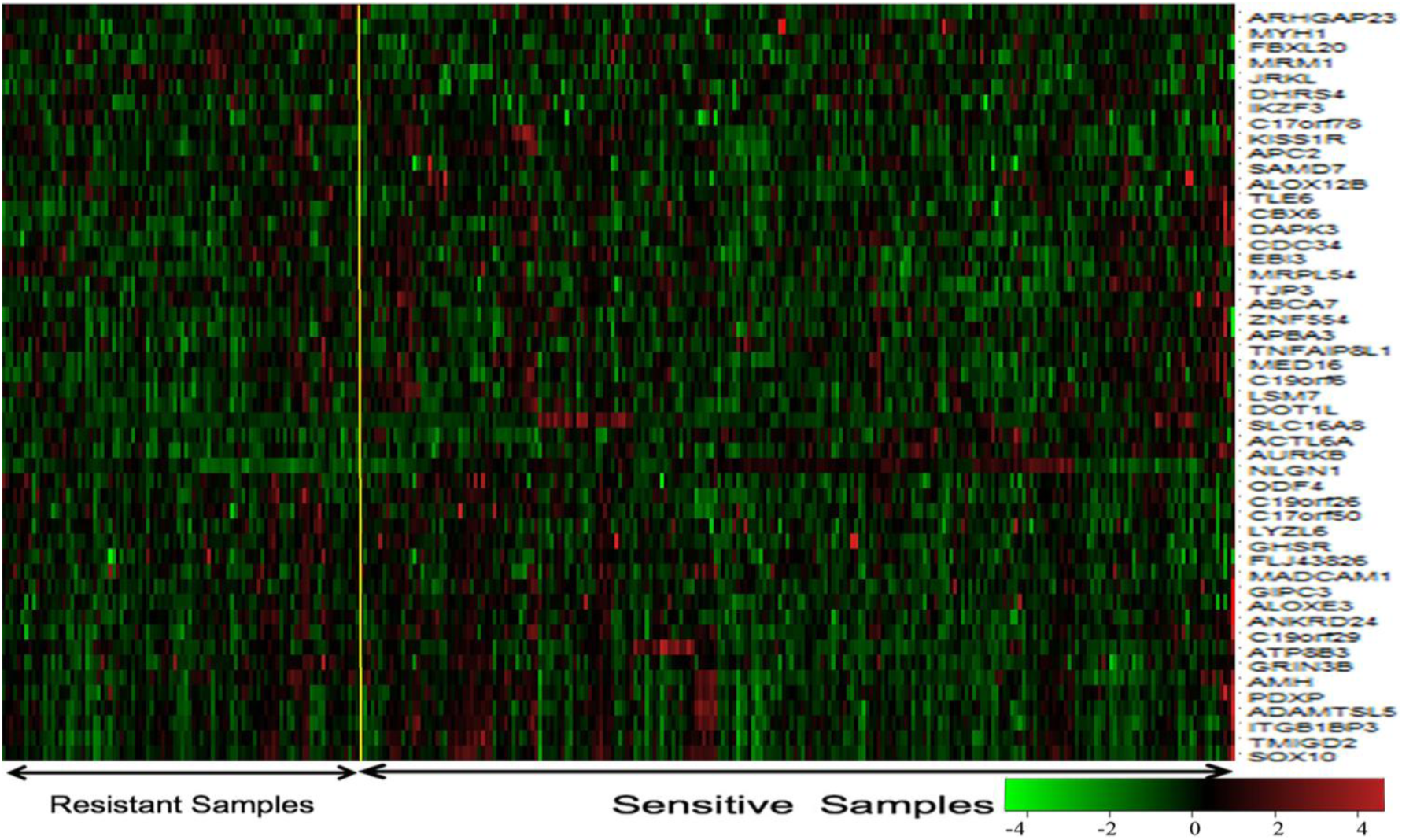
The gene expression heatmap of the top 50 candidate modulators.

### Dosage-sensitive genes and dosage-resistant genes and driver patterns

We conducted the module networks learning with the gene modules and candidate modulators. It generally assumed that the modulators most likely regulate the expression of the genes in the corresponding gene modules. Owing to the gene modulators would have distortions in amount of samples and they are likely to be the roots of the regression (drive) trees, the modulator gene are generally treat as a driver gene. Through the module network learning, we obtained 456 modules and 93 modulators (Additional file 7). 5 out of these modulators are from somatic mutations. Remarkably, several modules shared the same modulators, in which only one modulator is derived from somatic mutations.

We applied a local polynomial regression fitting model with the R package loess to the scatter diagram for 93 modulators with 324 samples. DS threshold is set to (-0.25, 0.25) as the filtering criteria for dosage-sensitive genes (DSGs) and the others were selected as dosage-resistant genes (DRGs). Therefore, we obtained 44 DSGs and 49 DRGs (Additional file 8). The LRG1, HYDIN, POLRMT and TNFSF10 gene data scatter diagram for CNVs or somatic mutations vs. gene expression with local polynomial regression fitting model are shown in Fig. 3, in which HYDIN is derived from somatic mutations.

**Figure 3.**
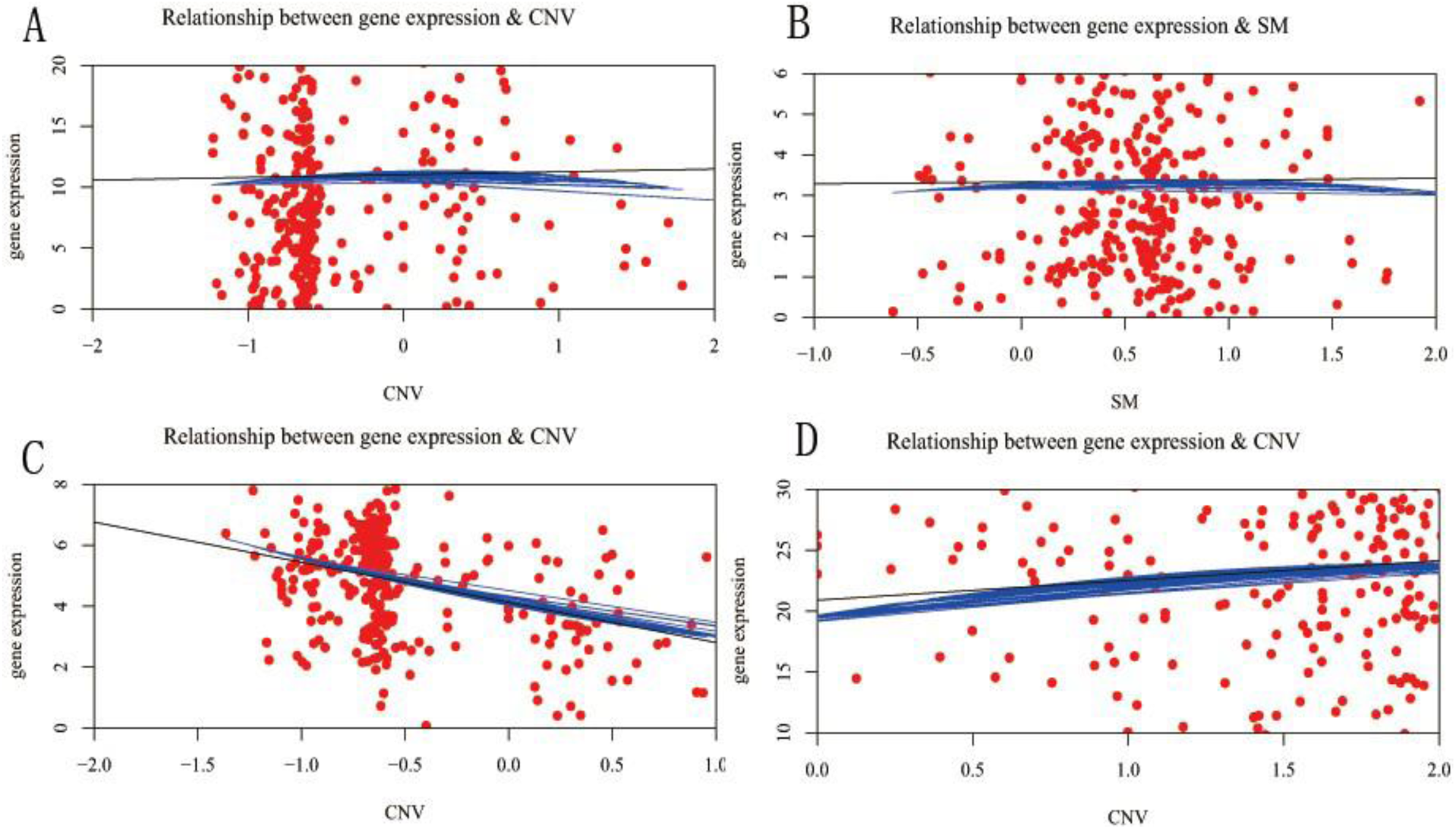
Gene data scatter diagram with local polynomial regression fitting model. Each red point represents an independent sample. The black line is the linear regression result, and the blue curve is the LOESS regression result. (A) LRG1. (B) HYDIN. (C) POLRMT. (D) TNFSF10.

Each red point represents an independent sample. The black line is acquired from the linear regression model, and the blue curve is resulted from the local polynomial regression fitting model. In Fig. 3A and Fig. 3B the slope of the black lines are close to 0, so the dosage sensitivity score are nearly equal to 0. In Fig. 3C and Fig. 3D the slope’s absolute value of the black line are above 1, the blue curves are mostly overlapped with the black line, so the dosage sensitivity score are above 1. Therefore, LRG1 and HYDIN are dosage-resistant genes, whereas POLRMT and TNFSF10 are dosage-sensitive genes.

We identified three major driver patterns in which the top 3 gene modules with the largest number of regulatory genes (modulators). Fig. 4 shows these driver patterns including gene modulators and the corresponding gene modules. Gene modulators were shown in the left side with blue background ovals are the dosage-resistant genes. Gene modulators are in the right side are dosage-sensitive genes. In the first driver pattern with color pale blue, there are 15 genes in gene co-expression module 3 and 13 gene modulators. In the second driver pattern with color green, there are 7 genes in module 30 and 18 gene modulators. In third pattern with color red, 6 genes are in module 31 and 14 gene modulators are for regulatory. Remarkably, it shows that CCL11 regulate all the 3 modules; SLC18A1 and C9orf82 regulate the module 3 and module 30; PLXDC1, CCL16, CDC34 and HYDIN regulate the module 3 and module 31; ZFHX4 regulates the module 30 and module 31. Hence, these important regulatory genes regulate many gene modules, and these gene modulators are highly correlated with ovarian cancer (Nolen et al. 2010; Willis et al. 2016; Priebe et al. 2008; Manabe et al. 2011; Kanska et al. 2016; Morrison et al. 2016; Chugh et al. 2016).

**Figure 4.**
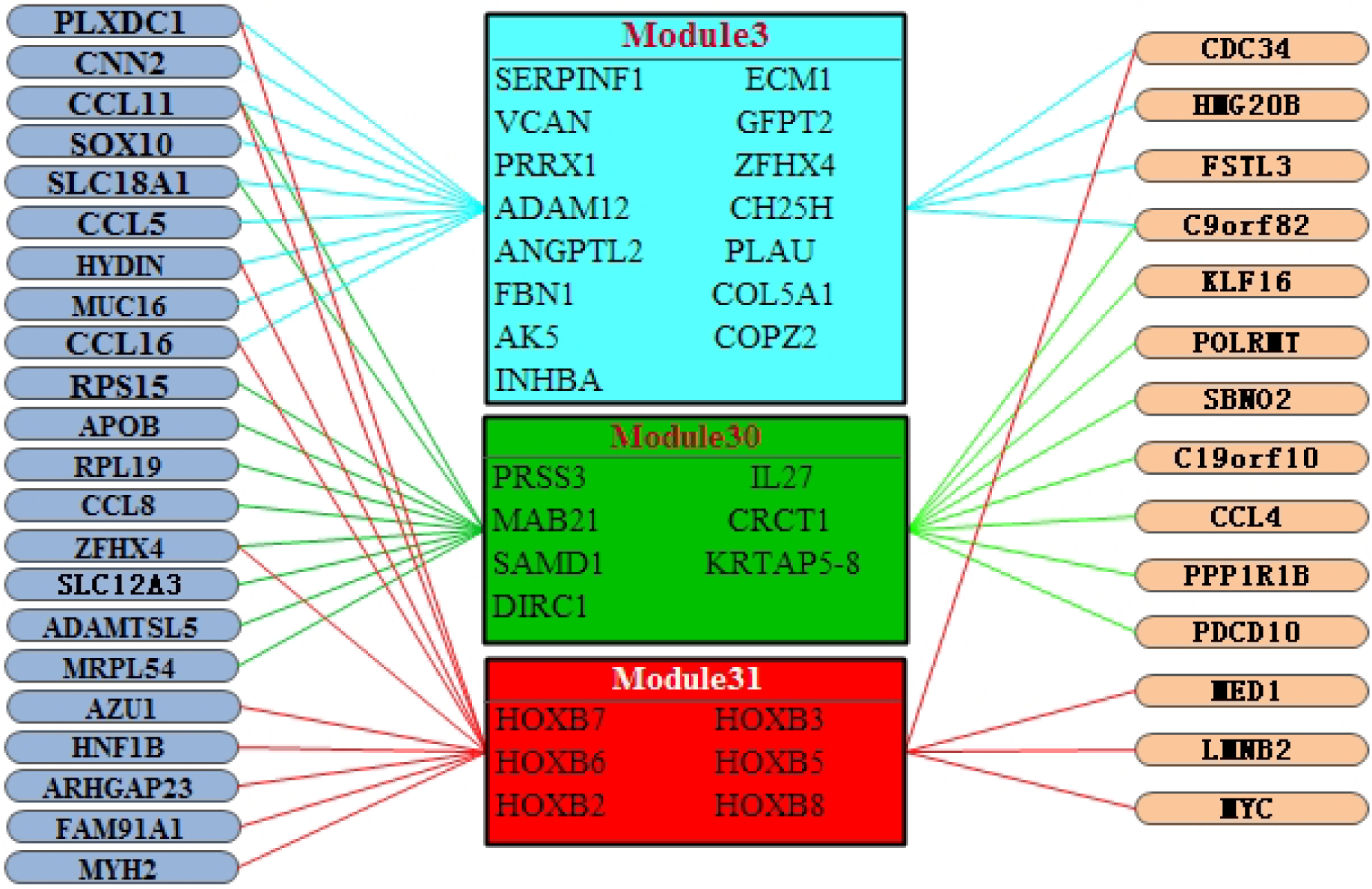
The 3 major driver patterns for dosage-sensitive and dosage-resistant. The graph depicts inferred regulators (left; ovals with blue background represents the DRGs, and ovals with white background represents the DSGs) and their corresponding regulated gene modules (right).

### Enrichment analysis for driver patterns

We applied the Gene Ontology (GO) category to expand the results of DRGs and DSGs with the R package clusterProfiler, in which the significance threshold are set as p-value < 0.05 and q-value < 0.05 separately. The top 12 enriched GO terms for DRGs and DSGs are presented in Table 2 and Table 3, respectively. The DRGs are mainly enriched in receptor binding, signaling pathway, cell migration, cellular response, cell chemotaxis, activator activity, etc. These processes are highly relevant to the trait of tumor and contribute to the chief progression of tumor. And the DSGs are mainly enriched in receptor binding, signaling pathway, template transcription, cell proliferation, cell differentiation, tissue development, cell division, etc. Compared with DRGs, the GO annotated to DSGs are mostly management functions and basic functions of the cell and tissue. Fig. 5 shows the top 12 biological process annotation analysis results on GO terms for DRGs and DSGs, respectively. The length of the bar represents the number of selected genes annotated onto this GO term, and the colors of the bar represents the significance levels of enrichment. The GO enrichment analysis results for DRGs and DSGs are in Additional file 9 and Additional file 10, respectively. From GO biological process enrichment analysis, we achieved CCL11, CCL16, CCL18, CCL23, CCL8, CCL5, APOB, BRCA1 play extremely significant role in DRGs, and EEF2, GNA11, GNA15, MED1, MYH8 and TCAP act very important role in DSGs. CCL11, CCL16, CCL18, CCL23, CCL8 and CCL5 have effective chemical attraction on eosinophil catalyzing their accumulation at allergic inflammation sites. CCL11 has been shown to negatively regulate neurogenesis with certain human tumors (Nolen et al. 2010; Ku et al. 2017). CCL16 is a chemokine prominently expressed in the liver, but also in ovarian and breast cancer (Manabe et al. 2011; Gantsev et al. 2013). CCL18 is associated with some human cancer types including ovarian cancer (Urquidi et al. 2012; Wang et al. 2015). CCL23 plays a role in subclinical systemic inflammation and associated with atherogenesis (Ignacio et al. 2016). CCL8 is strengthened in stromal fibroblasts at the tumor border and in tissues at which breast cancer cells incline to metastasize such as the lungs and the brain (Farmaki et al. 2016; Zsiros et al. 2015). CCL5 is a chemokine that boosts cancer progression by arousing and adjusting the inflammatory diseases, which sequently reconstruct the cancer microenvironment. Moreover, CCL5 also boost metastasis in ovarian and breast cancer cells (Zsiros et al. 2015; Soria et al. 2008). APOB mutation is associated with some human cancer types such as steatosis, liver, hypocholesterolemia (Cefalu et al. 2013; Williams et al. 2016). BRCA1 mutation confers high risks of ovarian and breast cancer, encodes a tumor suppressor (Rosenberg et al. 2016; Weren et al. 2016). EEF2 is a key components for protein synthesis, and ubiquitously expressed in normal cells (Kaul et al. 2011). GNA11 is associated with Congenital Hemangioma (Ayturk et al. 2016). GNA15 is associated with inhibition of proliferation, activation of apoptosis and differential effects (Zanini et al. 2015). MED1 is associated with some biological process such as prostate cancer cell growth (Liu et al. 2010). MYH8 is a key components for muscle development and regeneration (Tonami et al. 2013). TCAP is involved with energy regulation and metabolism, and is implicated in the regulation of stress-related behaviors (Aquila et al. 2016).

**Table 2.** The top 12 enriched GO terms for DRGs.

| GO-BP Term | pvalue | qvalue | gene |
| --- | --- | --- | --- |
| lymphocyte chemotaxis | 3.43E-09 | 2.12E-06 | CCL11/CCL16/CCL18/CCL23/CCL8/CCL5 |
| monocyte chemotaxis | 4.76E-09 | 2.12E-06 | CCL11/CCL16/CCL18/CCL23/CCL8/CCL5 |
| mononuclear cell migration | 1.95E-08 | 5.80E-06 | CCL11/CCL16/CCL18/CCL23/CCL8/CCL5 |
| chemokine-mediated signaling | 3.4E-08 | 5.87E-06 | CCL11/CCL16/CCL18/CCL23/CCL8/CCL5 |
| lymphocyte migration | 3.67E-08 | 5.87E-06 | CCL11/CCL16/CCL18/CCL23/CCL8/CCL5 |
| neutrophil chemotaxis | 3.95E-08 | 5.87E-06 | CCL11/CCL16/CCL18/CCL23/CCL8/CCL5 |
| cellular response to interleukin-1 | 4.91E-08 | 6.25E-06 | CCL11/CCL16/CCL18/CCL23/CCL8/CCL5 |
| neutrophil migration | 6.05E-08 | 6.25E-06 | CCL11/CCL16/CCL18/CCL23/CCL8/CCL5 |
| myeloid leukocyte migration | 6.31E-08 | 6.25E-06 | CCL11/CCL16/CCL18/CCL23/CCL8/CCL5/AZU1 |
| granulocyte chemotaxis | 1.09E-07 | 9.41E-06 | CCL11/CCL16/CCL18/CCL23/CCL8/CCL5 |
| cellular response to tumor | 1.16E-07 | 9.41E-06 | APOB/CCL11/CCL16/CCL18/CCL23/CCL8/CCL5/ BRCA1 |
| granulocyte migration | 1.84E-07 | 1.36E-05 | CCL11/CCL16/CCL18/CCL23/CCL8/CCL5 |

**Table 3.** The top 12 enriched GO terms for DSGs.

| GO-BP Term | pvalue | qvalue | gene |
| --- | --- | --- | --- |
| dopamine receptor signaling | 2.38E-06 | 0.001887 | PALM/GNA15/GNA11/KLF16 |
| skeletal muscle contraction | 0.000127 | 0.032678 | EEF2/TCAP/MYH8 |
| multicellular organismal movement | 0.000239 | 0.032678 | EEF2/TCAP/MYH8 |
| musculoskeletal movement | 0.000239 | 0.032678 | EEF2/TCAP/MYH8 |
| trabecula morphogenesis | 0.000255 | 0.032678 | MED1/SBNO2/UBE4B |
| DNA-templated transcription | 0.000289 | 0.032678 | MED1/MED16/POLR2E/MED24/POLRMT |
| phospholipase C-activating | 0.000289 | 0.032678 | GNA15/GNA11 |
| natural killer cell chemotaxis | 0.000352 | 0.03489 | CCL7/CCL4 |
| negative regulation of keratinocyte | 0.000498 | 0.038042 | MED1/KDF1 |
| androgen receptor signaling | 0.000544 | 0.038042 | MED1/MED16/MED24 |
| cardiac muscle tissue morphogenesis | 0.000544 | 0.038042 | MED1/TCAP/UBE4B |
| response to radiation | 0.000691 | 0.038042 | PPP1R1B/CCL7/GNA11/MYC/ECT2/UBE4B |

**Figure 5.**
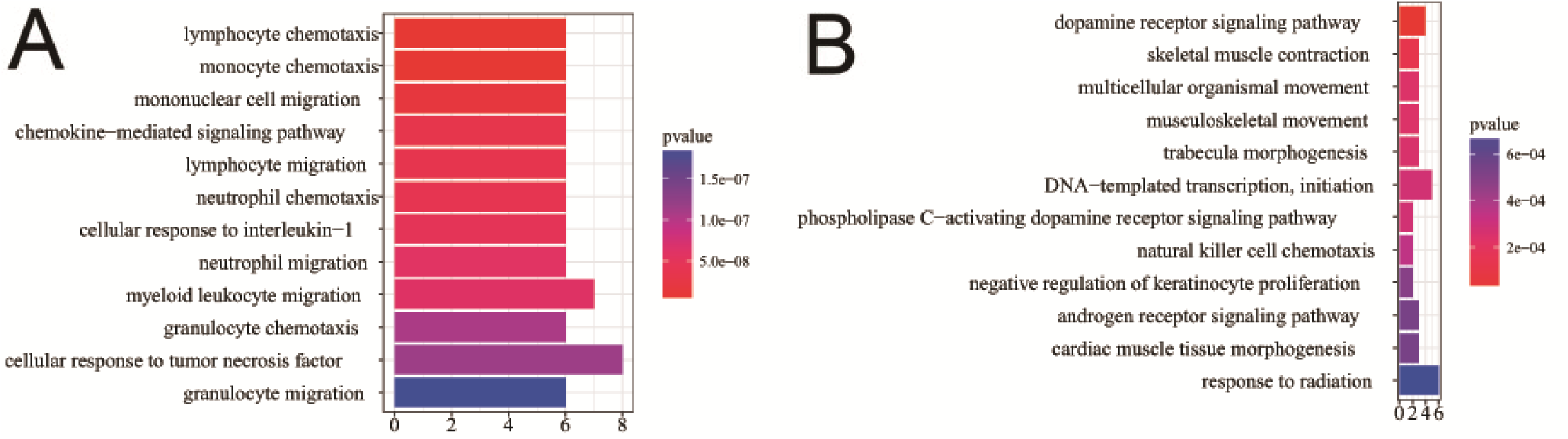
The analysis of GO biological process annotation GO biological process enrichment of DSGs analysis for DSGs and DRGs. (A)The GO biological process enrichment of DRGs. (B) The GO biological process enrichment of DSGs

### Regulatory pathway analysis for driver patterns

To further elucidate the biological pathway of gene modulators in the driver pattern, we performed the Kyoto Encyclopedia of Genes and Genomes (KEGG) pathway enrichment analysis of the results of DSGs and DRGs list with the R package clusterProfiler separately. The top 12 pathways for DRGs and DSGs are presented respectively in Table 4 and Table 5. For DRGs, the most significant canonical pathways are mainly bound up with Chemokine signaling, Cytokine-cytokine receptor interaction, Parkinson’s disease, Breast cancer, onset diabetes, Asthma, Prion diseases, etc. And the pathways for DSGs are major related to Thyroid hormone signaling, Apoptosis, Cytosolic DNA-sensing, Amoebiasis, Chagas disease, Cell cycle, WNT signaling, etc. Remarkably for DRGs, Cytokine-cytokine receptor interaction is embedded in the Chemokine signaling pathway. For these two pathways, gene CCL11, CCL16, CCL18, CCL23, CCL8 and CCL5 have a key function (see Fig. 6). Chemokines play a role in the migration of many cells during development and are critical to nervous system development (Li et al. 2004). Cytokine interactions play an important role in health and are vital to many cancer during immunological and inflammatory responses in disease. It can lead to antagonist, additive, or synergistic activities in keeping physiological functions such as body temperature, feeding and somnus, as well as in anorectic, ardent fever, and sleepiness neurological representations of acute and chronic disease (Lippitz et al. 2013). Parkinson’s disease is a pathway specifically associated to diseases of the central nervous system, NDUFB5 and SLC18A1 are enriched in this pathway (see Fig. 7). NDUFB5 and SLC18A1 are associated with multiple human diseases (Willis et al. 2016; Chen et al. 2014). Breast cancer is the major cause of cancer death among the female worldwide, gene BRCA1 and FGF22 are enriched in this pathway (see Fig. 8). FGF22 is required for brain development and associated with hereditary and neoplastic disease (Heroult et al. 2014). For DRGs, the thyroid hormones are key regulators of metabolism, growth and other body systems, MED1, MED16, MED24 and MYC are play a role in this pathway (see Fig. 9). MYC is involved in cell cycle progression, apoptosis, and cellular transformation and is related to many tumors (Kim et al. 2015). Apoptosis is an evolutionarily conserved signaling pathway, which plays a fundamental role in regulating cell number and eliminating damaged or redundant cells (Zeestraten et al. 2013), GADD45B, TNFSF10, LMNB2 are enriched in this pathway (see Fig. 10). TNFSF10 is associated with some human cancer types including ovarian cancer (Charbonneau et al. 2013). LMNB2 acts as regulators of cell proliferation and differentiation, regards as a cancer risk biomarker in several cancer subtypes (Sakthivel et al. 2016). Cell cycle progression is completed through a reproducible sequence of events. It is a significant pathway associated with many diseases including ovarian cancer (Taylor_Harding et al. 2013). GADD45B and MYC are enriched in this pathway (see Fig. 11). GADD45B is implicated in some responses to cell injury including cell cycle checkpoints, apoptosis and DNA repair (Salvador et al. 2013). WNT signaling pathway is required for basic developmental processes and play an important role in human stem cells and cancers (Krausova et al. 2014). MYC and TBL1XR1 are enriched in this pathway (see Fig. 12). The KEGG pathway enrichment analysis result for DRGs and DSGs are shown in Additional file 11 and 12, respectively.

**Table 4.** The top 12 enriched GO terms for DSGs.

| pathway | pvalue | gene |
| --- | --- | --- |
| Chemokine signaling pathway | 4.80E-06 | CCL11/CCL16/CCL18/CCL23/CCL8/CCL5 |
| Cytokine-cytokine receptor | 4.58E-05 | CCL11/CCL16/CCL18/CCL23/CCL8/CCL5 |
| Parkinson's disease | 0.052537 | NDUFB5/SLC18A1 |
| Breast cancer | 0.053866 | BRCA1/FGF22 |
| Other types of O-glycan biosynthesis | 0.056251 | MFNG |
| Ribosome | 0.060691 | RPS15/RPL19 |
| Vitamin digestion and absorption | 0.061213 | APOB |
| Maturity onset diabetes of the young | 0.066151 | HNF1B |
| NOD-like receptor signaling | 0.070724 | MFN1/CCL5 |
| Phototransduction | 0.071064 | RCVRN |
| Asthma | 0.078387 | CCL11 |
| Prion diseases | 0.088067 | CCL5 |

**Table 5.** The top 12 enriched GO terms for DSGs.

| pathway | pvalue | gene |
| --- | --- | --- |
| Thyroid hormone signaling | 0.000438 | MED1/MED16/MED24/MYC |
| Apoptosis | 0.009077 | GADD45B/TNFSF10/LMNB2 |
| Cytosolic DNA-sensing pathway | 0.017271 | POLR2E/CCL4 |
| NF-kappa B signaling pathway | 0.036048 | GADD45B/CCL4 |
| Amoebiasis | 0.036744 | GNA15/GNA11 |
| Chagas disease | 0.041034 | GNA15/GNA11 |
| Cholinergic synapse | 0.048583 | PIK3R5/GNA11 |
| Cytokine-cytokine receptor | 0.05237 | CCL7/CCL4/TNFSF10 |
| Cell cycle | 0.058258 | GADD45B/MYC |
| FoxO signaling pathway | 0.065054 | GADD45B/TNFSF10 |
| Ubiquitin mediated proteolysis | 0.069434 | CDC34/UBE4B |
| Wnt signaling pathway | 0.074816 | MYC/TBL1XR1 |

**Figure 6.**
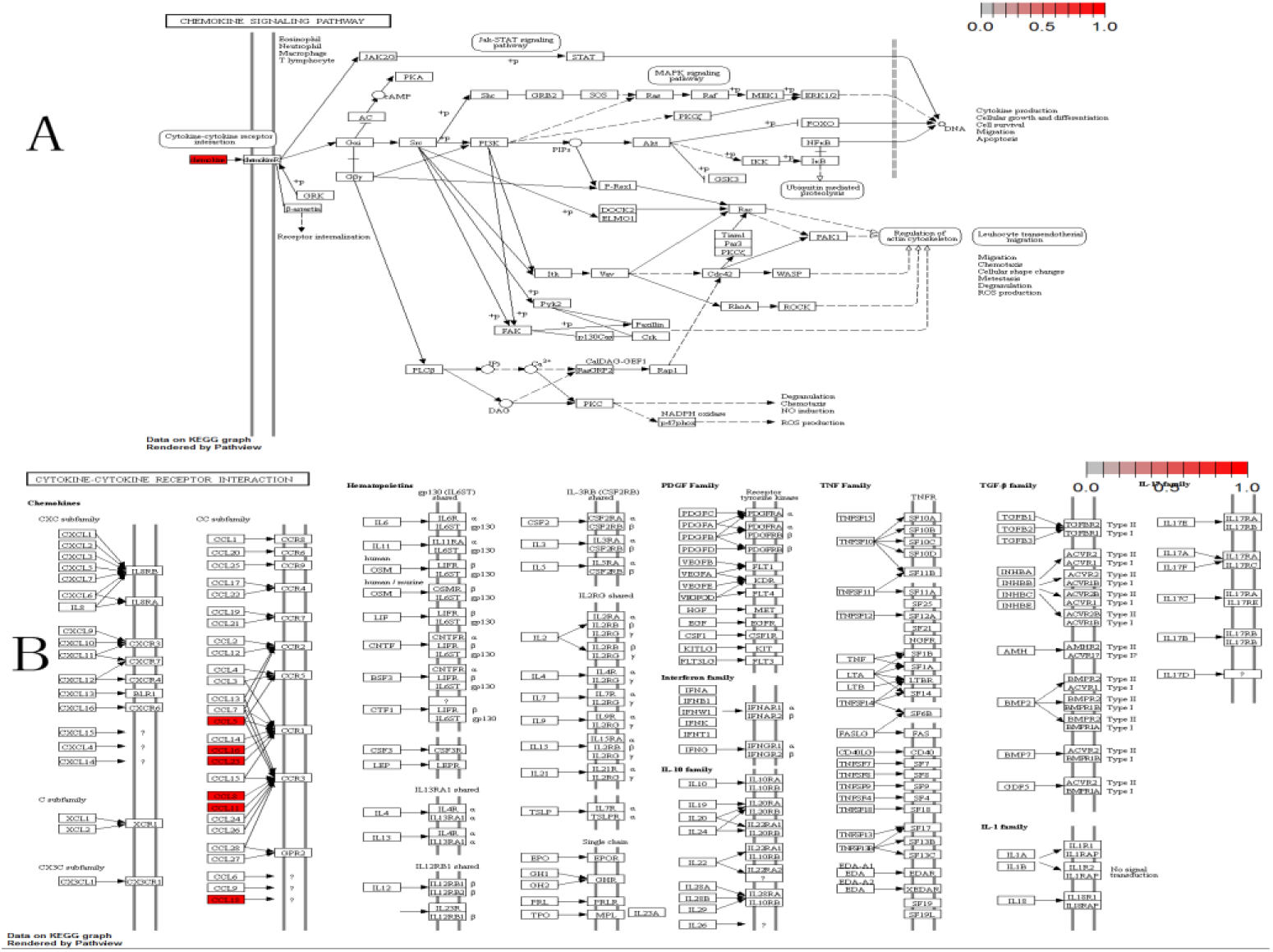
The six genes are enriched in the top 2 significant pathways (red, up-regulated). (A) Chemokine signaling pathway. (B) Cytokine-cytokine receptor interaction.

**Figure 7.**
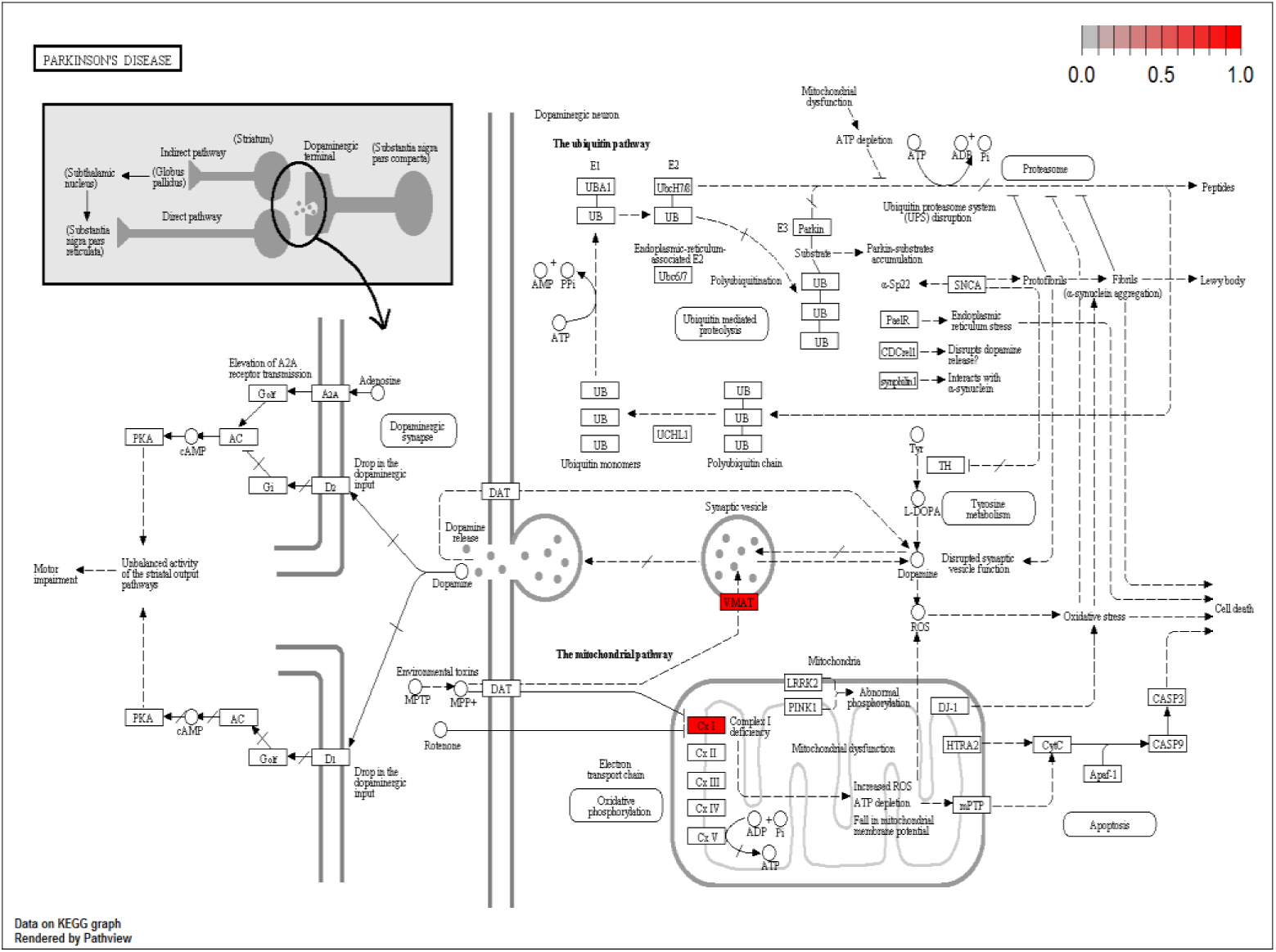
Parkinson disease pathway(red, up-regulated). NDUFB5 and SLC18A1 are enriched in this pathway. NDUFB5 is a gene alias of the complex 1 deficiency(Cx1) and SLC18A1 is a gene alias of the vesicular amine transporter(VMAT).

**Figure 8.**
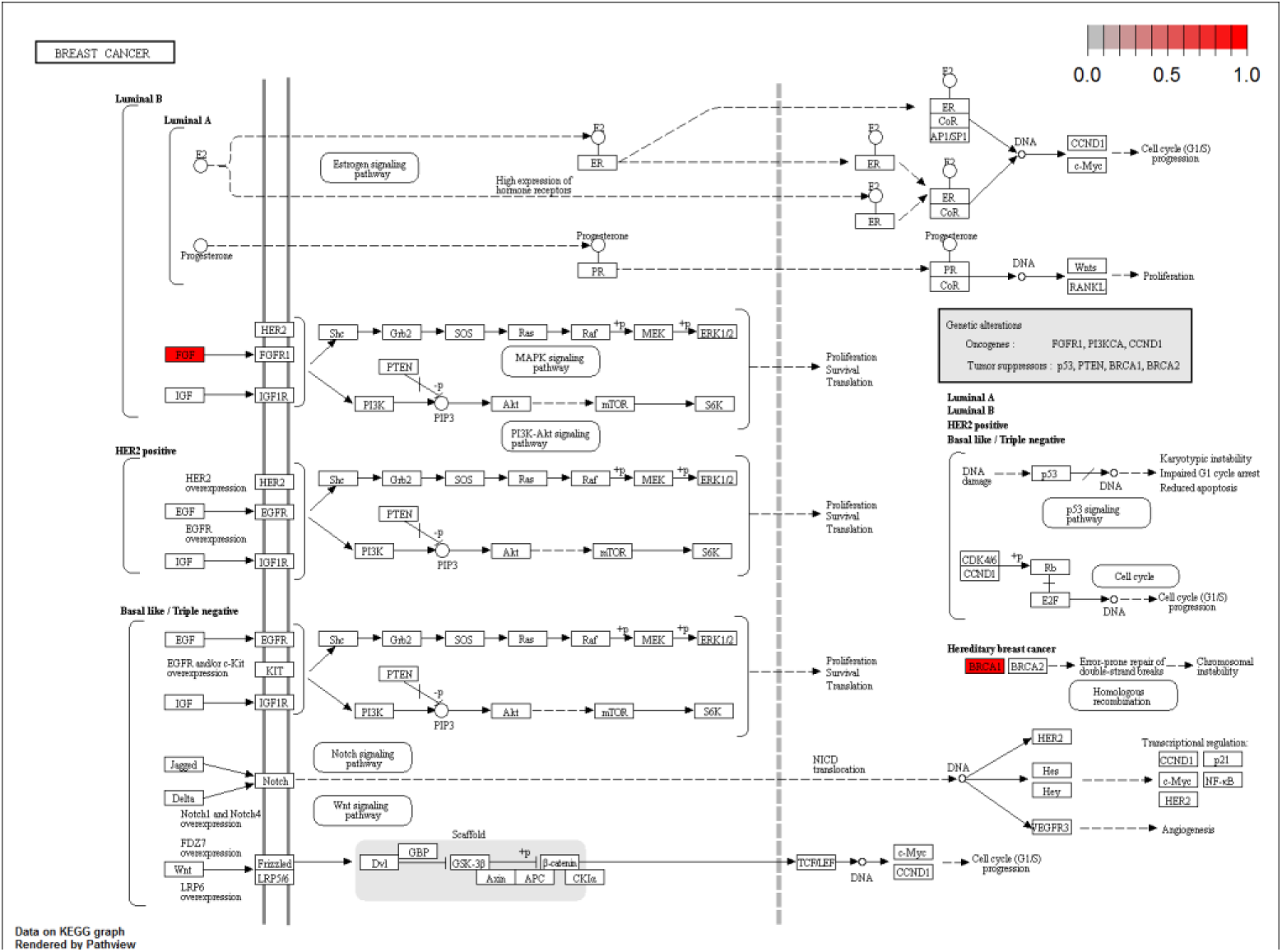
Breast cancer pathway (red, up-regulated). BRCA1 and FGF22 are enriched in this pathway.

**Figure 9.**
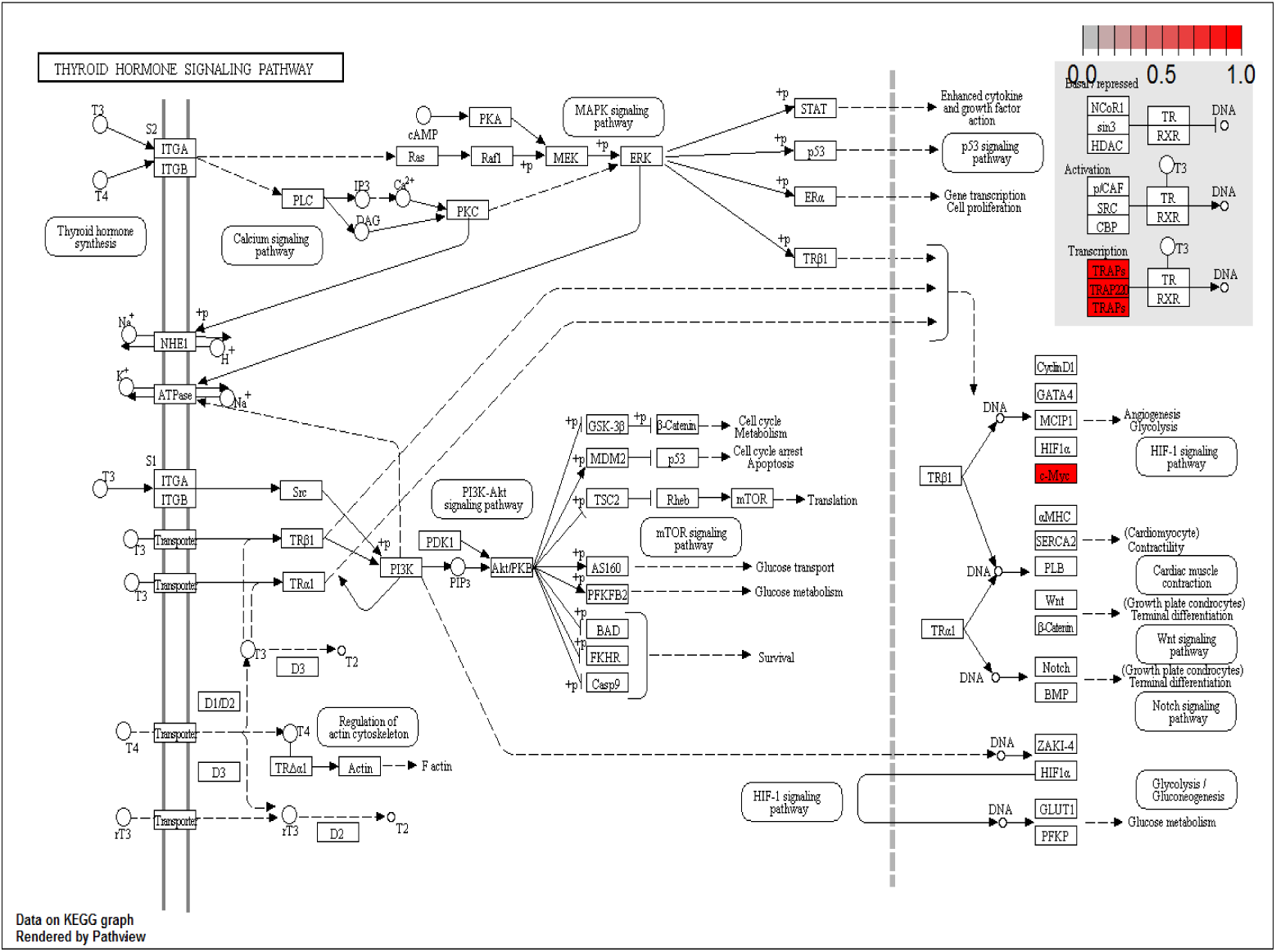
thyroid hormones pathway (red, up-regulated). MED1, MED16, MED24 and MYC are enriched in this pathway.

**Figure 10.**
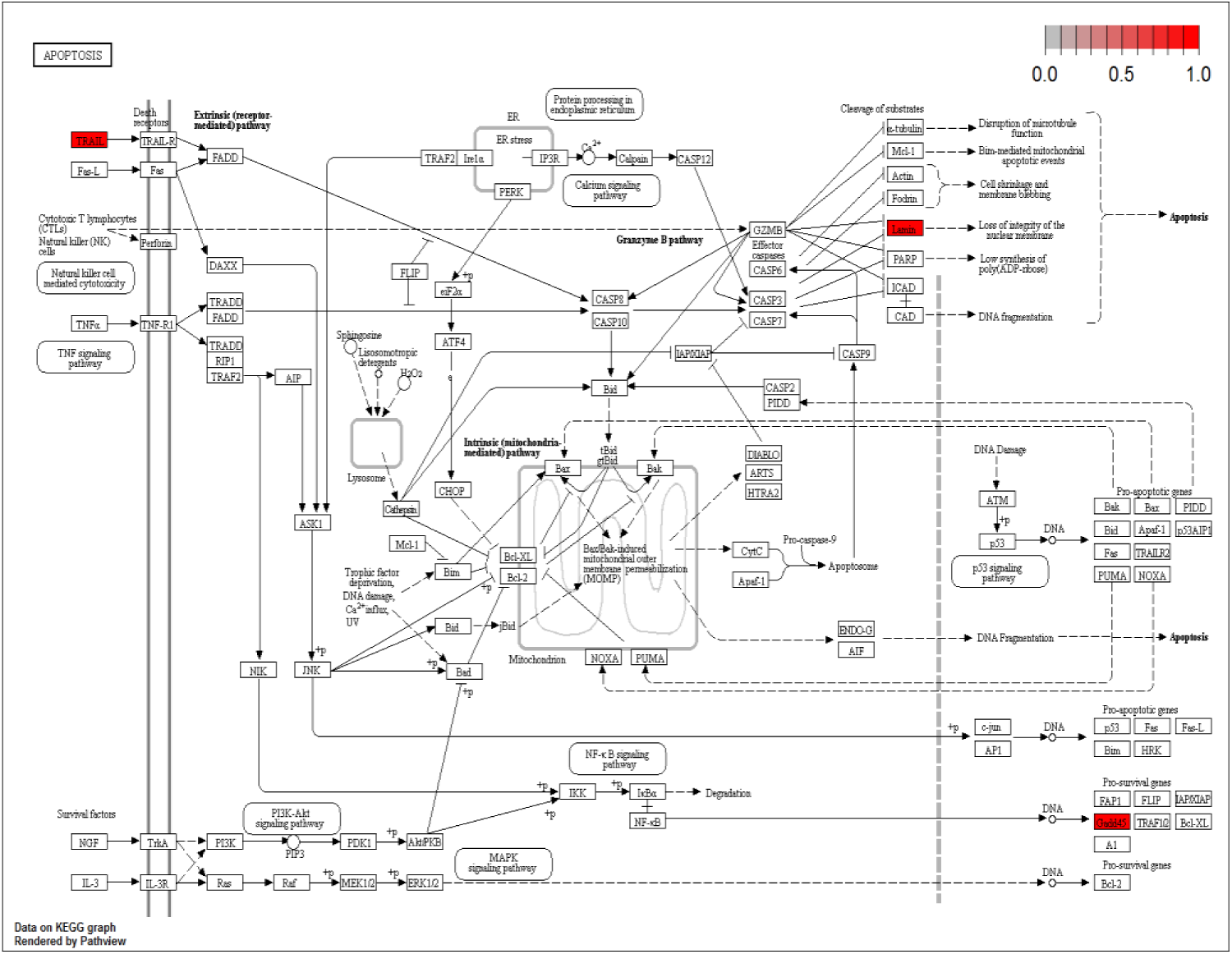
Apoptosis pathway (red, up-regulated). GADD45B, TNFSF10, LMNB2 are enriched in this pathway. TRAIL is a gene alias of TNFSF10, LMNB2 is a gene alias of the lamin.

**Figure 11.**
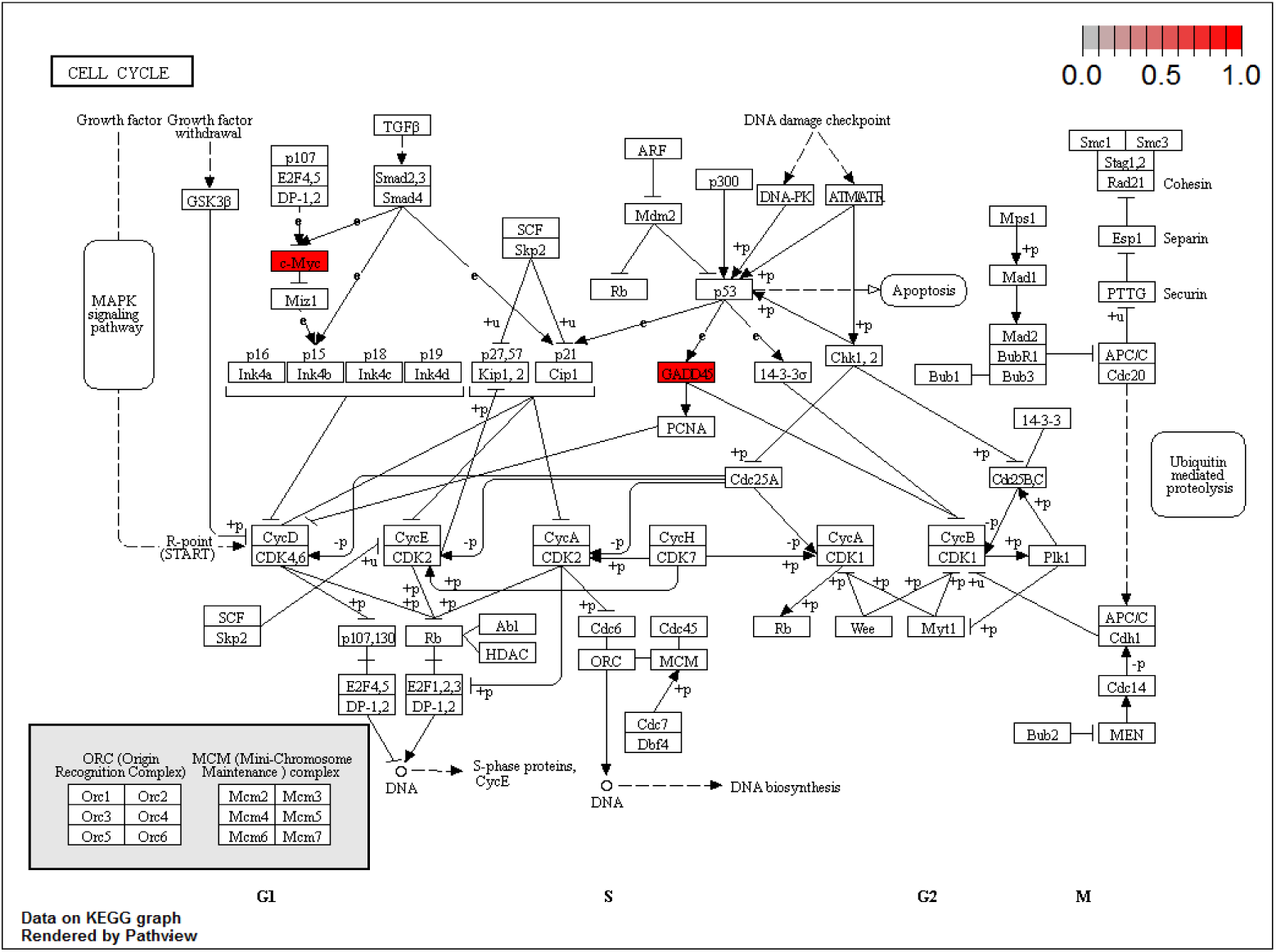
Cell cycle pathway (red, up-regulated). GADD45B and MYC are enriched in this pathway.

**Figure 12.**
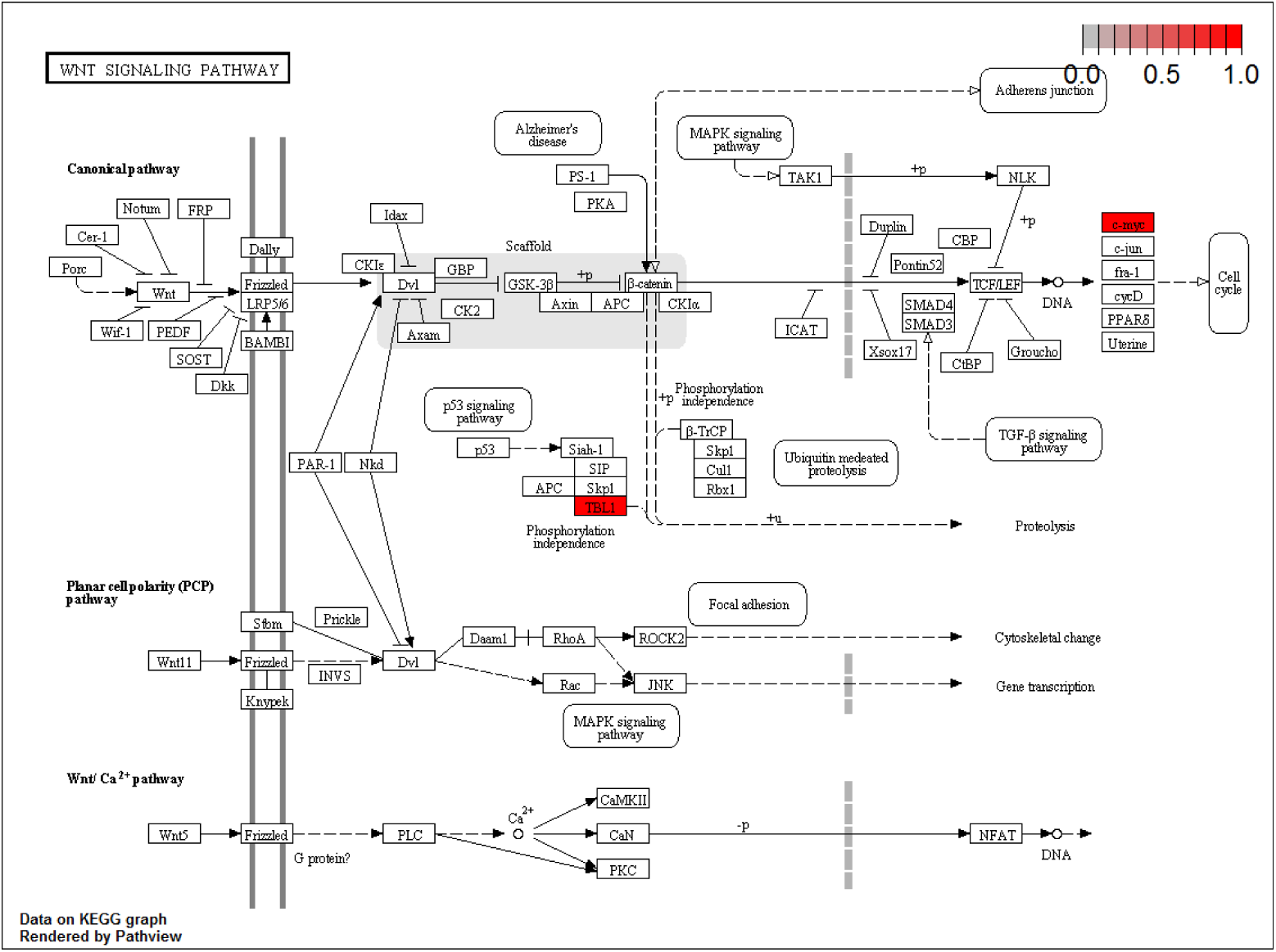
Wnt signaling pathway (red, up-regulated). MYC and TBL1XR1 are enriched in this pathway.

## Conclusions

The main purpose of this research was to distinguish the candidate driver genes and the corresponding driving mechanism for resistant and sensitive tumor from the heterogeneous data. We presented a machine-learning approach to integrate somatic mutations, CNVs and gene expression profiles to distinguish interactions and regulations for dosage-sensitive and dosage-resistant genes of ovarian cancer. We developed a general framework for integrating high dimensional heterogeneous omics data and can potentially lead to new insight into many related studies at the systems level. In the framework, weighted correlation network analysis (WGCNA)is applied to the co-expression network analysis; the mutation network is constructed by integrating the CNVs and somatic mutations and the initial candidate modulators is selected from the clustering the vertex of network; the regression tree model is utilized for module networks learning in which the obtained gene modules and candidate modulators are trained for the modulators regulatory mechanism; finally, a local polynomial regression fitting model is applied to identify dosage-sensitive and dosage-resistant driver patterns. From the Gene Ontology and pathway enrichment analysis, we obtained some biologically meaningful gene modulators, such as CCL11, CCL16, CCL18, CCL23, CCL8, CCL5, APOB, BRCA1, SLC18A1, FGF22, GADD45B, GNA15, GNA11 and so on, which can be conducive to appropriate diagnosis and treatment to cancer patients.

## Methods

We integrated gene expression, CNV, and somatic mutation data to identify the driver patterns for resistant and sensitive tumors. The overview of the overall integrative analysis is shown in Fig. 1. As can be seen in Fig. 1, the procedure takes these datasets as inputs and then produces a short list of genes as candidate divers. The method contains the following steps: (1) extract a pool of candidate modulators as initial driver genes, which have significant CNVs/Somatic mutations in ovarian cancer samples and achieve the initial gene modules from the gene expression by weighted correlation network analysis (WGCNA); (2) construct the heterogeneous network by module networks learning and evaluate the regulatory mechanism between these different omics data by liner regression model; (3) identify driver patterns and modulators interactions in the constructed regulatory networks using gene ontology and pathway enrichment analysis.

**Figure 13.**
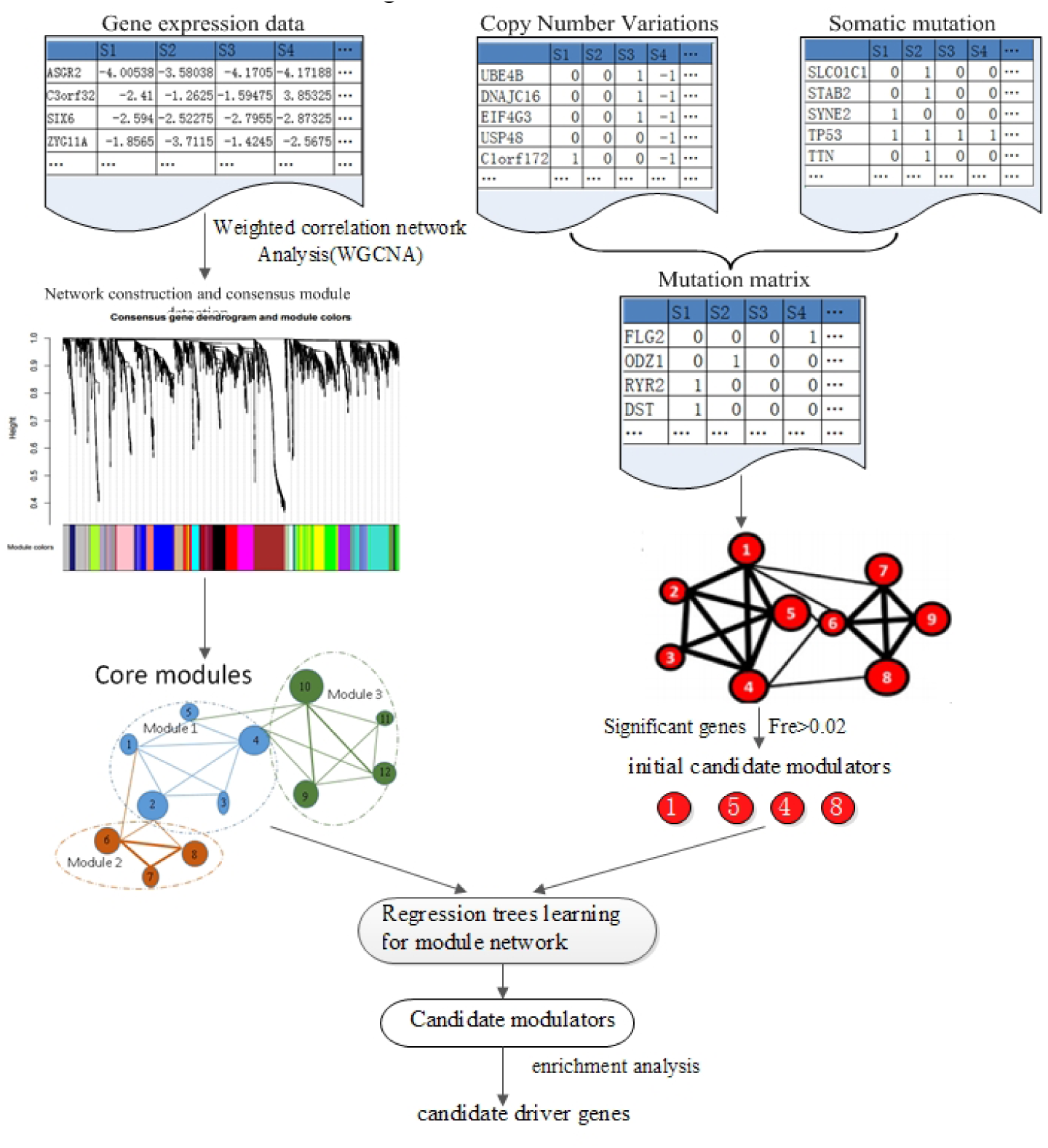
the scheme of driver patterns identification. acquire the initial gene modules for gene expression profiles via WGCNA network analysis (on the left)and extract a pool of candidate modulators as initial driver genes integrate CNVs and Somatic mutations in ovarian cancer samples (on the right); (2) construct the heterogeneous network by module networks learning and evaluate the regulatory relationships by liner regression model;(3) identify modulators interactions in the constructed regulatory networks and analyze the driver patterns through and gene ontology analysis and pathway enrichment analysis.

### Resources and datasets

We downloaded gene expression data from the Agilent 244 K Custom Gene Expression platform, CNV data from the Affymatrix Genome-Wide Human SNP Array 6.0 platform, and somatic mutation data from whole exome sequencing data obtained using Illumina Genome Analyzer DNA Sequencing. The clinical data of these patients are analyzed to determine cis-platinum chemotherapy response samples and these tumor samples are classified into resistant and sensitive groups, sensitive tumors have a platinum-free space of six months or more after the last primary treatment, no sign of residue or relapse, and the follow-up will at least six months later; and resistant tumors are recurred within six months after the last treatment (Prado_2014). According to the above definition, we identified 93 platinum-resistant and 231 platinum-sensitive primary tumors from the TCGA website, for which the gene expression profiles, CNV, and somatic mutation data were available as well. The 324 samples list are shown in Additional file 1. The gene expression data are listed in Additional file 2. The CNVs data are listed in Additional file 3. The somatically mutated genes are listed in Additional file 4.

### Gene co-expression modules analysis using WGCNA

Weighted correlation network analysis is a framework for co-expression analysis to explore gene modules of high correlation to external sample traits. The gene expression correlation networks are based on quantitative measurements correlations. Given an *n* × *m* expression matrix X where the row indices correspond to genes (*i* = 1, …,*n*) and the (*i* =1, …,) column indices correspond to samples:

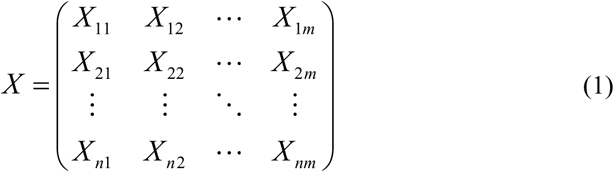

The co-expression similarity *S_ij_* is applied to calculate the adjacency matrix:

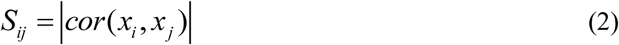

Where *S_ij_* is the absolute value of the correlation coefficient between gene i (*i* = 1, …, *n)* and gene j *(j* = 1, …,*n*) in the expression profiles matrix X. The value of *S_ij_* is in [0, 1]. The adjacency function *a_ij_* is defined by soft threshold of co-expression similarity *S_ij_* by soft threshold *β* as following:

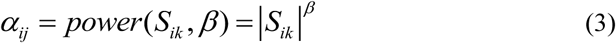

Where *β>1.* To measure the topological structure with adjacency function, topological overlap matrix (TOM) is applied to build the matrix *Ω=[ω_ij_]*,

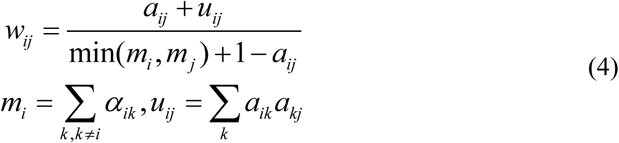

In which the network structure is not only based on the two genes with direct adjacency correlation, but also on the indirect adjacency relationship.

The gene expression network is constructed based on matrix Ω. Gene modules are detected by unsupervised hierarchical clustering method. Give the qth gene co-expression module, the eigengene expression is *E^(q)^*. The correlation coefficient for the gene *x_i_* and the module *E^(q)^* can be defined as:

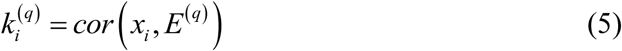

Where 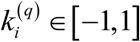

### Modulators identification using module networks learning

From the initial gene co-expression modules, we focused on the candidate modulators’ regulatory by regression tree model. Each candidate modulator is connected to a group of genes by maximizing the normal-gamma scoring function which describes the qualitative behavior of a candidate modulator that regulate the gene expression modules. A regression tree is formed with two basic blocks: decision nodes and leaf nodes. Each node corresponds to one of the selected modulators above. Our learning is an iterative process. In each iteration, a regulation procedure is found for each gene module and then each gene reassigned to the corresponding module due to prediction with the best regulatory function. We searched for the model with the highest score by using the Expectation Maximization (EM) algorithm. The scoring function defined as:

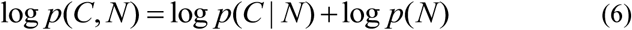

Where C is the candidate data and N the model structure of the network. The first term is the candidate data for a given model which meets normal gamma distribution. The second part is a penalty score on network complexity. The EM algorithm ensures the reliability of the model until convergence to a local maximum score. The essence of the algorithm consists of two steps: the first procedure is learning the best regulation program (regression tree) for each module. The tree is constructed from the root to its leaves. The second procedure is to find the association rules for corresponding module.

### Dosage-sensitive and dosage-resistant analysis

From the module networks learning, the candidate genes were generated. Then we applied a local polynomial regression fitting model to conduct dosage-sensitive and dosage-resistant analysis. In the model, a predict function was adopted to obtain n isometric points on the LOESS curve (Yan_2017). To achieve the monotonicity of the copy number expression interrelation a function is defined as M:

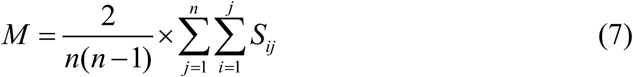

In which

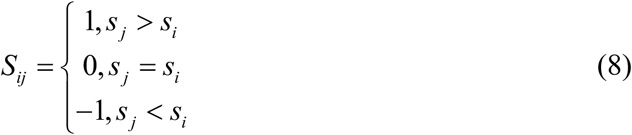

Where n is the number of samples. To quantize the association between gene expression and CNVs (somatic mutations), another linear fitting model was adopted and a straight line with slope of K was obtained. The slope of linear model can reflect the relationship of variables in LOSS model to some extent. Hence the dosage sensitivity (DS) score can be defined as:

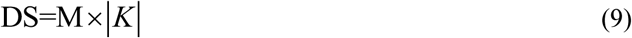

Herein, for a gene, a larger DS indicates stronger dosage sensitivity.

### Data access

The CNV, mutation and gene expression data are used in this paper can be downloaded from Broad GDAC Firehose website (http://gdac.broadinstitute.org/runs/analyses__2014_10_17/data/OV/20141017).

## Acknowledgements

The authors are grateful to Peter Langfelder’s team who provided the weighted correlation network analysis method with WGCNA package, Prof Ripley B. D. who provided the Local Polynomial Regression Fitting analysis method with LOESS package, Dana Pe’er Lab who provided the Module Networks learning analysis method with CONEXIC Toolkit and Guangchuang Yu’s team who provided the Functional enrichment analysis method with clusterProfiler package. This work was supported by the National Natural Science Foundation grant of China and China Scholarship Council grant.

